# 3DBrainOne: an integrated end-to-end platform for 3D histological analysis of whole mouse brains

**DOI:** 10.64898/2026.05.06.723327

**Authors:** Daye Kim, Young-Gyun Park

## Abstract

Three-dimensional (3D) whole-organ imaging and analysis at cellular resolution (termed 3D histology) provide profound insights into the organization and interactions of cells throughout organs. However, the quantitative analysis of these massive datasets remains a significant bottleneck due to the lack of integrated, user-friendly tools. Here, we present 3DBrainOne, an end-to-end ImageJ plugin that streamlines the essential 3D histological analysis of the mouse brain—from raw image preprocessing to region-wise quantification—within a single platform. 3DBrainOne features a robust whole-brain cell-counting module that uses a Difference-of-Gaussians (DoG) blob detection algorithm followed by a ResNet18-based deep learning classifier, enabling high-fidelity automatic whole-brain cell counting with a graphical user interface (GUI) for visual inspection and manual curation of analysis results. 3DBrainOne also supports multi-channel colocalization analysis. Furthermore, this platform includes modules for atlas alignment and brain-region-wise volumetric quantification, enabling brain region-resolved cell counting and structural analyses. As an ImageJ plugin, 3DBrainOne is compatible with a range of operating systems and hardware. In summary, 3DBrainOne is an integrated, versatile, and easy-to-use platform that will facilitate 3D histological analyses in experimental neuroscience.

## Main

In recent years, there has been a growing demand in neuroscience for three-dimensional (3D) whole-brain imaging and quantitative analyses [1–5]. This trend reflects the recognition of the limitations inherent in conventional two-dimensional (2D) imaging, including the loss of 3D spatial context, difficulties in assessing interregional interactions, low sensitivity and throughput for screening cellular and molecular features, and sampling bias [6]. In a typical 2D histological study, only thin tissue sections (<100um in thickness) are examined, which can obscure the continuity of structures and neural circuits across slices and introduce sampling bias. By contrast, 3D imaging provides continuous structural information and 3D spatial context, enabling a more comprehensive and unbiased understanding of brain organization and functions, highlighting a systems-level perspective and the distributed nature of neural information processing [7].

At the same time, single-cell-level analyses are indispensable, as cells are the fundamental units and compartments of biological functions. Accurate assessment of intercellular relationships, molecular identities, and spatial topologies can be achieved via single-cell analyses. When integrated with region-wise volumetric analysis, these high-resolution datasets provide clues on how individual cells may contribute to the physiology and functions of the brain.

While advances in 3D histological techniques (e.g., tissue clearing, organ-scale immunolabeling, and light-sheet fluorescence microscopy) enable whole-organ labeling and imaging [1, 6, 8], software tools capable of quantitatively analyzing organ-scale image datasets at single-cell resolution remain limited. Most widely used image analysis platforms, such as ImageJ/Fiji [9], CellProfiler [10], and QuPath [11] are primarily optimized for 2D data. Software tools for 3D histological analyses, such as ClearMap [12], cellfinder [26], brainreg [13], and Neuroglancer [14] have enabled biological discoveries. However, their practical utility as comprehensive, user-friendly analysis platforms remains limited, as they provide only a subset of essential 3D histological analyses (e.g., cell counting, atlas alignment, brain-region-resolved quantification) or require complex configuration, scripting, or command-line operations, making them less accessible to users without a computational background. Consequently, quantitative image analysis remains a major bottleneck in 3D histological studies.

We present 3DBrainOne, an all-in-one graphical user interface (GUI)-based plugin of ImageJ [9] for automated 3D histological analysis of the mouse brain **(Fig. 1)**. The platform enables automatic, accurate cell detection and atlas registration of whole-mouse brain images, with a GUI for visual investigation and manual correction of the data, enabling accurate mapping of cells and volumes at the brain-region level. The cell counting module incorporates a supervised model pre-trained on whole-brain datasets acquired under diverse imaging conditions, allowing direct application to new datasets without additional labeling. The platform further supports multi-channel co-positivity analysis to identify cells co-positive for multiple marker signals. In parallel, its atlas-aligning module enables brain region-resolved analysis of detected cells and, when combined with the volumetric quantification module, enables brain region volume comparison across brains. To enhance usability, 3DBrainOne allows visual inspection and parameter adjustment by overlaying detection and classification results directly on the image data. In addition, its post-processing functionalities, including region-wise normalization and manual correction of detected blobs, improve the reliability and interpretability of the results.

**Fig. 1.**
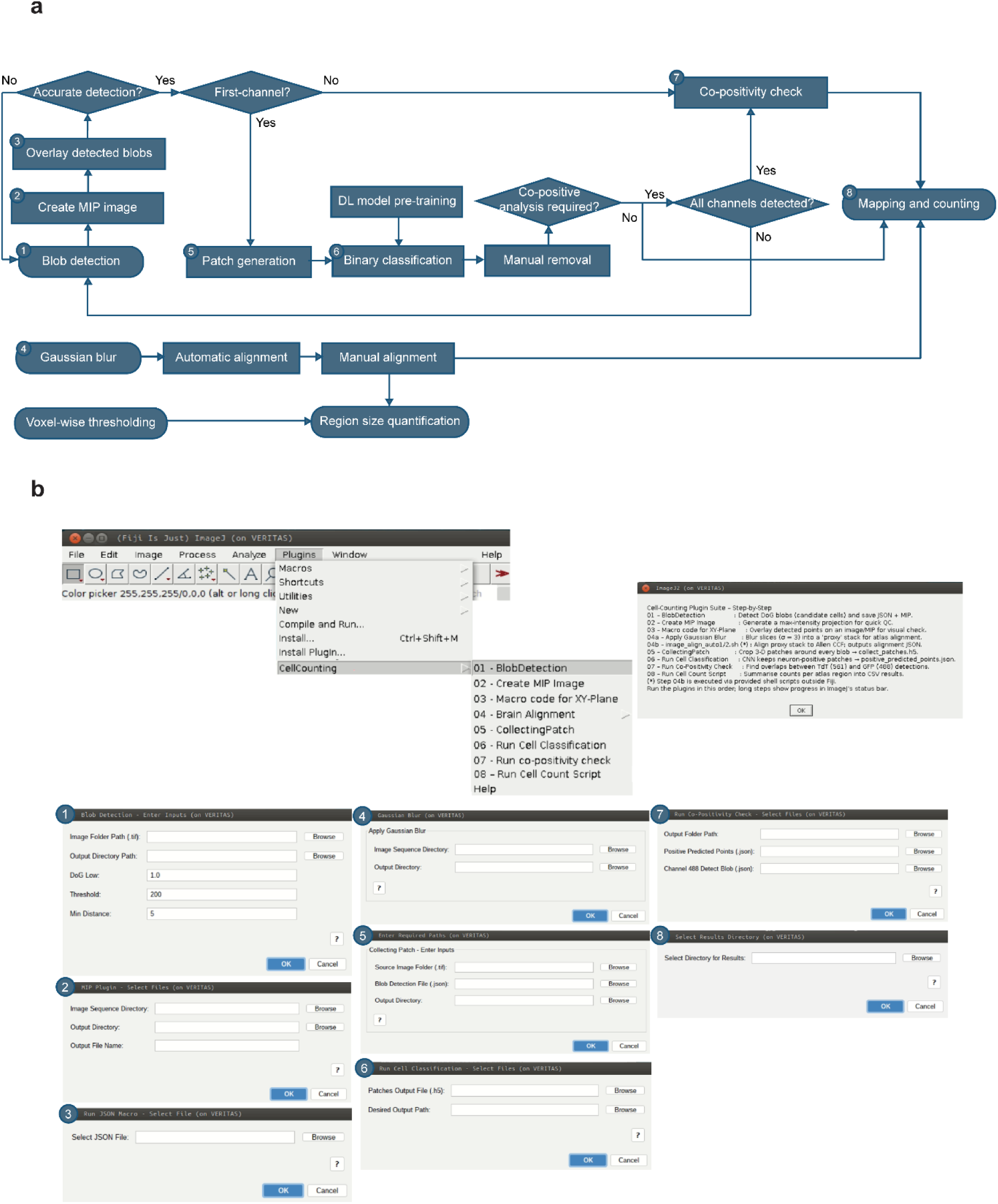
Overview of the 3DBrainOne platform and plugin interface. **(a)** Schematic diagram of the 3DBrainOne analysis platform. The workflow comprises sequential modules for cell counting, atlas alignment, and volumetric quantification of brain regions. **(b)** User interface of the 3DBrainOne ImageJ plugin. Windows for the individual steps of **a** process are numbered. Dedicated input windows allow parameter tuning (e.g., DoG sigma, threshold, patch size), and a built-in help panel guides users through the recommended execution sequence.

## Results

### 3DBrainOne enables end-to-end whole-brain image analysis

3DBrainOne enables detection of genetically labeled individual cells across whole mouse brain images, followed by atlas registration and region-wise quantification **(Fig. 1a)**. The platform is implemented as a GUI-based ImageJ plugin **(Fig. 1b)** that automates the entire workflow within a unified, user-accessible platform [9].

The analysis proceeds through a series of integrated modules. Cell candidates are initially identified using a Difference-of-Gaussians (DoG)-based blob detector [15], followed by multi-view patch extraction to capture local image features across orthogonal planes. These candidates are subsequently refined using a deep learning–based classifier built on a ResNet18 architecture [16], with optional user-defined threshold fine-tuning to adapt to varying signal conditions. The model was validated across 21 mouse whole-brain datasets spanning diverse imaging conditions and signal characteristics.

The platform further incorporates co-positivity analysis to identify cells with overlapping signals across fluorescent channels, and anatomical registration to the Allen Mouse Brain Atlas (CCFv3) [17] for standardized spatial mapping. Based on this registration, region-wise quantification is performed to compute cell counts and densities, accompanied by post-processing and visualization steps to ensure data interpretability.

The annotation of detected cells can be overlaid directly on whole-brain images, allowing users to interactively set thresholding parameters, review detected cell distributions, and manually remove suspected false positives within the GUI. The result can be exported in multiple formats, including CSV and heatmap formats.

To showcase 3D histological analysis of whole mouse brains using 3DBrainOne, we analyzed Ai14 mouse brains injected with Cre-expressing adeno-associated virus (AAV). Ai14 mice [18] are genetically engineered reporter lines in which the expression of the fluorescent protein tdTomato is driven by Cre recombinase activity. We used a viral construct that expresses Cre recombinase and enhanced green fluorescent protein (eGFP) fusion protein via neuron-specific human synapsin (hSyn) promoter [20].

### Stage 1. Whole-Brain Imaging Data Acquisition and Preprocessing

Following a two-weeks of AAV injection into the Motor cortex of the Ai14 mouse brains, we processed the mouse brains using SHIELD, a tissue protection-based tissue clearing technique [21]. After transcardially perfusing the mice with SHIELD-perfusion solution, the brains were harvested and post-fixed in the solution at 4°C for 48 hours. Then the brain was passively delipidated in a buffer containing 300 mM sodium dodecyl sulfate, 100 mM sodium sulfite, and 10 mM boric acid (pH 9.0) at 45 °C for 12–21 days. Following extensive washing to remove residual detergents, optical clearing was performed by equilibrating the tissues in an optical clearing solution (OCS; 2,2’-thiodiethanol, dimethyl sulfoxide, and iohexol-based solution) until full transparency was achieved. The optically cleared samples were mounted in 1.5% (w/v) agarose in OCS and incubated in an OCS-filled chamber overnight to ensure equilibration before imaging.

Whole brains were imaged in their intact form using a light-sheet microscope (SmartSPIM, LifeCanvas Technologies, MA), equipped with a 3.6X objective lens (NA = 0.2; 1.8 µm/pixel) at excitation wavelengths of 488 nm (for visualizing eGFP), 561 nm (for visualizing tdTomato), and 647nm (for visualizing autofluorescence). Acquired image data were preprocessed using SmartSPIM software. Preprocessing steps included Z-stack alignment, stripe artifact removal (de-striping), and tile stitching, resulting in high-quality 3D image volumes suitable for downstream analysis. All datasets were standardized to a consistent voxel size to ensure compatibility across samples and facilitate automated processing and quantitative comparison. The final preprocessed 3D brain volumes served as input to the subsequent analysis stages.

### Stage 2. Cell Candidate Detection

The preprocessed 3D whole-brain image data consisted of two fluorescent channels: 561 nm (tdTomato) and 488 nm (eGFP), which were analyzed independently. For each channel, cell candidates were detected using a blob-detection algorithm based on Gaussian DoG filtering and intensity thresholding. This approach identifies local maxima corresponding to potential cells by detecting localized intensity peaks [15].

Three key parameters were central to the detection process: the lower sigma value for DoG filtering (--dog-low), the minimum distance between detected peaks (--min-distance), and the intensity threshold (--threshold). These parameters were optimized individually for each sample, considering variability in fluorescent signal strength, tissue density, and background noise. To guide this optimization, maximum intensity projection (MIP) images were generated along the horizontal (xy) plane for each volume. Detected blob coordinates were overlaid on the MIP, and threshold settings were iteratively adjusted based on visual inspection. By comparing detection outcomes across threshold values with anatomical features visible in the MIP, we minimized false positives and false negatives, ensuring optimal sensitivity and specificity per channel **(Fig. 2a)**.

**Fig. 2.**
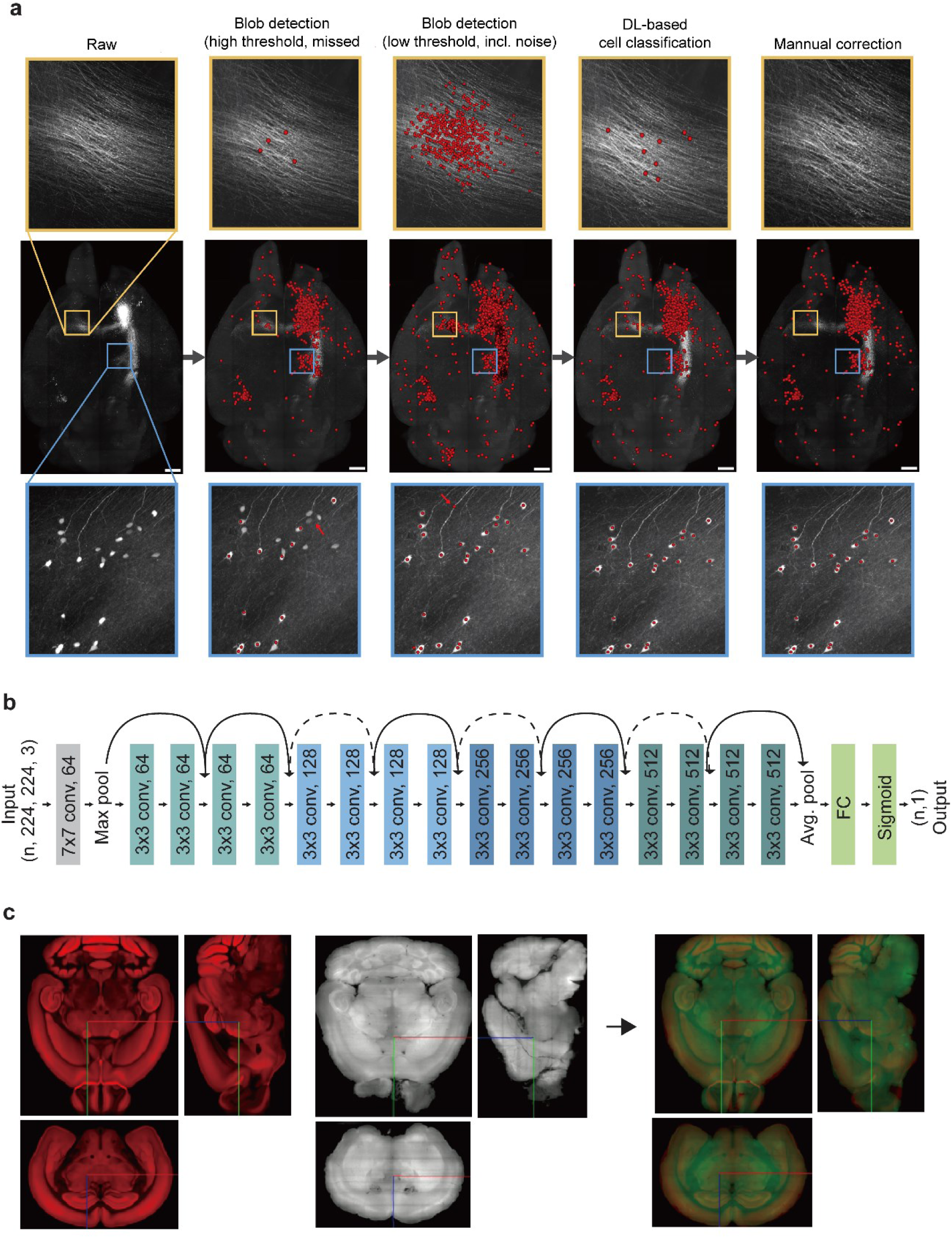
Representative images illustrating the cell counting module. **(a)** Representative images illustrating threshold tuning and classifier-based refinement. Blob detection was performed over a range of threshold values, and an optimal threshold was selected by visual inspection to maximize inclusion of true cells, even at the expense of minor noise. A deep learning–based classifier was subsequently applied to filter out false positives. Final results could be manually curated to ensure accuracy. Shown are representative outputs from raw image input, threshold-based blob detection, classifier output, and optional manual correction. **(b)** An architecture of the ResNet18-based binary classifier. Each detected blob candidate is represented by 2D image patches from three orthogonal planes (xy, yz, xz) and classified using a convolutional neural network trained on uniformly sampled brain-wide data. The classifier filters out false positives, thereby improving the precision of cell detection. **(c)** Atlas registration and anatomical localization. Light-sheet volumes are registered to the Allen Mouse Brain Atlas using both automated and manual alignment steps. Final cell coordinates are mapped into standardized 3D brain space, enabling region-wise quantification and visualization across samples.

For the tdTomato channel (561 nm), which typically exhibited strong and well-defined fluorescence signals, a relatively high threshold was applied to suppress background artifacts. In contrast, the eGFP channel (488 nm) often displayed weaker fluorescence, necessitating a lower threshold to preserve sensitivity and avoid omitting true positives. Importantly, this detection module is not limited to these channels and can be extended to other labeling modalities. By appropriately tuning parameters such as intensity threshold, distance, and Gaussian smoothing (sigma), the module may support detection of cells labeled with other markers such as DAPI, pending validation for each specific staining condition. The final output of this step was a list of blob center coordinates saved in JSON format. These coordinates served as input to the subsequent patch-generation and classification stages. At this point, the detected candidates had not yet been confirmed as true cells; this refinement was performed in the following step using a deep learning-based classifier.

### Stage 3. Patch Generation and Uniform Whole-Brain Labeling

For each cell candidate detected in the previous step, three 2D image patches (31×31 pixels) were extracted from the *xy*, *yz*, and *xz* planes, centered on the blob coordinate. This multi-view patch-generation strategy preserves the 3D structural context of the candidate region rather than relying solely on local intensity information from a single plane. The three orthogonal patches corresponding to each candidate were used jointly as input to the classification model to determine whether the candidate represented a true cell.

To construct a training dataset that uniformly represents the entire brain volume, we employed a structured sampling strategy. The whole-brain 3D image was divided into a 5×5×5 spatial grid, resulting in 125 subregions. From each subregion, a fixed proportion of candidate patches was randomly sampled to ensure spatial uniformity. This approach prevents overfitting to high-density regions and promotes robust performance in sparser brain areas, which are often underrepresented in naïvely sampled datasets.

Patch labeling was performed using a custom GUI designed for rapid manual annotation. Each patch was visually reviewed, and labels such as “Positive” or “Negative” were assigned using simple keyboard shortcuts. This efficient labeling workflow enabled rapid, high-quality training data at scale.

By encouraging spatial diversity and anatomical heterogeneity in the training dataset, this labeling strategy helps the model to avoid over-reliance on region-specific morphological features or signal intensities. As a result, the model achieves better generalization performance when applied to new samples or imaging conditions with varying anatomical and fluorescent characteristics.

### Stage 4. Deep Learning-Based Cell Classification

To determine whether each candidate blob corresponds to a true genetically labeled cell, we implemented a binary classification model based on a Residual Network (ResNet) architecture [16]. Specifically, we adopted the relatively lightweight ResNet18 variant, which is well-suited for this task due to its low parameter count, fast convergence, and proven reliability across a wide range of image classification problems. Its architectural simplicity and robust performance align well with the characteristics of our dataset **(Fig. 2b)**.

The model input is a 31×31×3 tensor, where each channel corresponds to a 2D image patch extracted from the *xy*, *yz*, and *xz* planes, respectively. The output is a binary classification (Positive vs. Negative) generated via a sigmoid activation function. Training was optimized using the binary cross-entropy loss function.

The training setup was configured as follows: optimizer: Adam (learning rate = 1e-4); batch size = 64; number of epochs = 50; data augmentation = 8× (including random rotations, flips, and Gaussian noise); validation split = 20%; cross-validation = 5-fold cross-validation.

Labeled patches used for training and testing were uniformly sampled from 21 whole-brain mouse samples, ensuring diverse imaging conditions and cell densities and supporting high generalization performance. All training procedures were conducted in a PyTorch environment [25], and overfitting was mitigated using early stopping based on validation loss.

### Stage 5. Inference and Post-processing

The trained classification model can be directly applied to new mouse brain samples without additional labeling. For each sample, the previously generated multi-view patches are input into the model, which automatically predicts whether each detected blob corresponds to a true cell. This step serves as a post-filtering mechanism that refines the initial threshold-based blob detection results. To further improve adaptability across datasets, users can adjust the decision threshold applied to the model’s probabilistic outputs, thereby providing flexible control over the precision–recall trade-off under varying signal-to-noise ratios (SNRs).

By leveraging the classifier to eliminate false positives from the over-detected candidate pool, the model achieves high precision while minimizing false negatives—an essential balance for robust cell detection in both densely and sparsely populated regions. The coordinates of blobs classified as positive (i.e., true cells) are saved in JSON format and serve as the input for the subsequent co-positivity analysis and region-wise quantification stages.

### Stage 6. Co-positivity Analysis

Based on the final cell coordinates independently detected and classified in the two fluorescent channels, we performed co-positivity analysis by comparing their spatial relationships. Specifically, Euclidean distances were computed between all positive cells across the two channels. When a cell from one channel was found within a predefined distance threshold of a cell from the other channel, the pair was considered co-positive—i.e., representing a single biological cell expressing both fluorescent signals.

The default threshold was set to 5 pixels, but the user could manually adjust it based on imaging resolution, expected cell size, and the biological characteristics of the cell types labeled in each channel. To determine the optimal threshold, co-localization results were visually overlaid and inspected to ensure that the classification of co-positive cells reflected true biological signal overlap.

Importantly, co-positivity analysis goes beyond detecting mere fluorescent overlap. It enables the identification of individual cells that simultaneously express both markers, thereby providing a quantitative basis for identifying cells expressing multiple markers. The spatial coordinates of co-positive cells were subsequently passed to the quantification module, where they were incorporated into region-wise analyses and directly compared against single-channel positive cell counts to enable multi-dimensional biological interpretation.

### Stage 7. Anatomical Registration to the Atlas

To assess the anatomical distribution of detected cells, we performed spatial registration of the preprocessed whole-brain images and associated cell coordinate data to the Allen Mouse Brain Common Coordinate Framework version 3 (CCFv3) [17] using nuggt (Chung lab; https://github.com/chunglabmit/nuggt). This registration process combined automated alignment with manual refinement, enabling accurate anatomical mapping and facilitating quantitative comparisons across multiple samples **(Fig. 2c)** [25].

For initial alignment, we generated MIP images from every 15th z-slice of the 488 nm channel, which contain autofluorescence signals reflecting the overall brain structure. These MIPs were rescaled to match the dimensions of the reference atlas and aligned using the SITK Align toolkit [22]. This step estimated affine and nonlinear transformations based on morphological similarity, including curvature, contours, and tissue boundaries. Landmark points were manually defined for each brain to guide and, when necessary, constrain the transformation process.

Following automated registration, the resulting alignments were visually inspected and manually refined in Neuroglancer, a web-based 3D image viewer [14]. This manual correction step was particularly important in regions with variable signal intensity or structural distortion, where automated alignment alone was insufficient. Final registration parameters were validated in all three planes to ensure anatomically meaningful region assignments and high spatial accuracy across samples.

### Stage 8. Quantification, Post-processing, and Visualization

Upon completion of atlas registration, each cell’s center coordinate was mapped to the corresponding region ID defined in the CCFv3 atlas [17], allowing for the automatic assignment of anatomical labels. Based on this mapping, the numbers of positive and co-positive cells in each anatomical region were computed, followed by downstream post-processing and visualization.

To ensure comparability across regions of different sizes, raw cell counts were normalized by the pre-defined area (or volume) of each region to yield cell density values. This normalization step provides a more accurate representation of relative cellular abundance by accounting for variations in region size.

The resolution of analysis was adjustable, either by selecting one of the hierarchical levels (1–7) of the Allen Brain Atlas or by applying user-defined anatomical masks. This enables the interpretation of results at varying spatial scales—from global whole-brain comparisons to detailed assessments of specific subregions.

Final quantification results were exported in CSV format, containing comprehensive information for each anatomical region, including region name, region ID, cell count, density, and region volume. These outputs were further processed using Python-based scripts to generate visualizations.

To improve analysis accuracy, users can manually remove false positives by selecting and deleting suspect cell coordinates directly in the image interface. This user-driven manual correction capability complements automated analysis, enhancing the reliability of the final results. All modifications were immediately reflected in the quantification data and subsequent statistical visualization.

### Robust cell detection across heterogeneous brain regions and imaging conditions

The performance of the developed cell classification model was quantitatively evaluated using a portion of the patch dataset collected from 21 whole-brain mouse samples—encompassing diverse imaging conditions and brain regions—as a held-out test set, using a 70:30 train-to-test split at the patch level, ensuring that no patches from the same spatial subregion were overrepresented in either set. The model was trained to perform binary classification between positive and negative classes, and performance was assessed using standard metrics derived from the confusion matrix **(Fig. 3a)** and the receiver operating characteristic (ROC) curve [23] **(Fig. 3b)**.

**Fig. 3.**
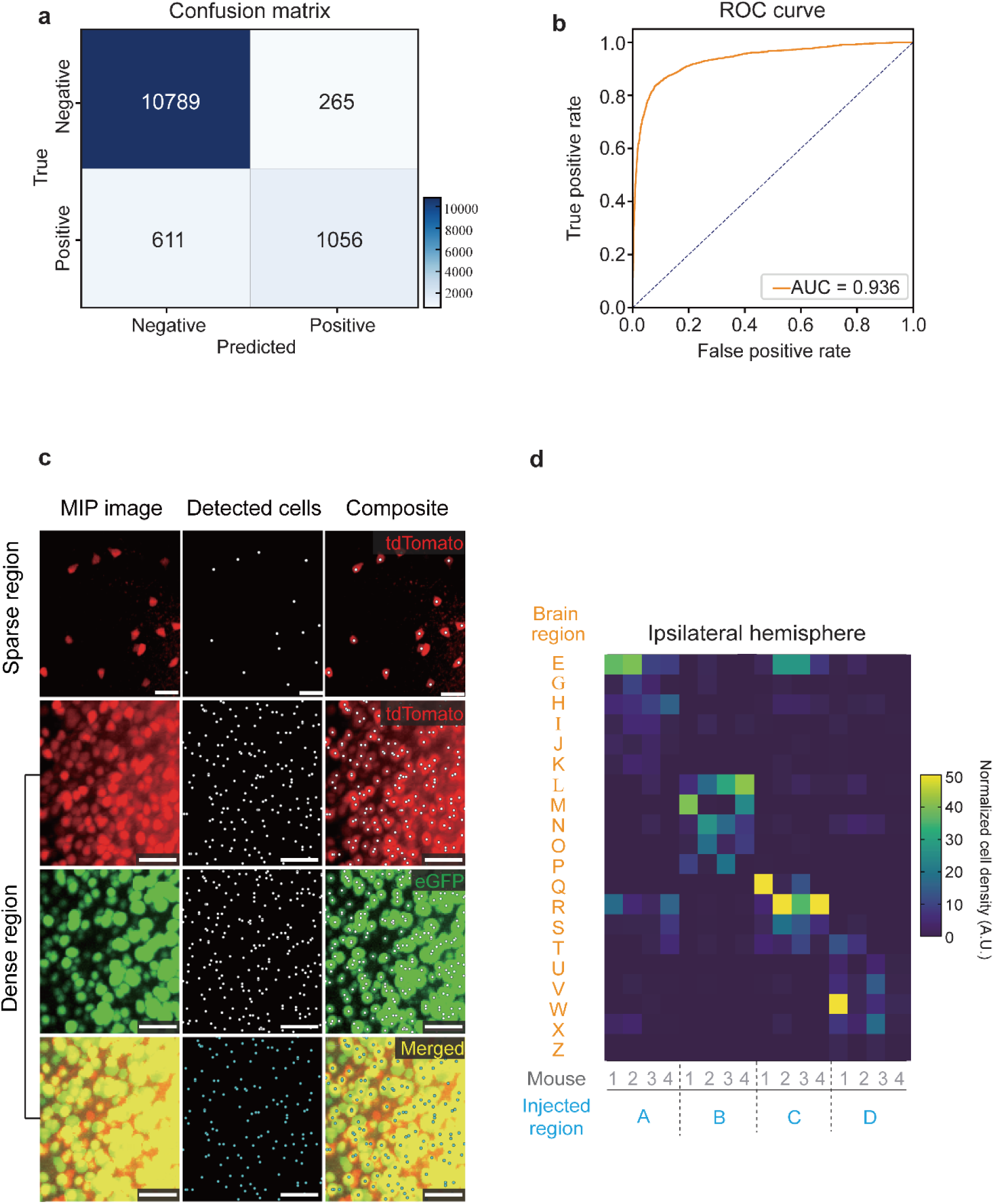
Classification performance and final cell detection results using the 3DBrainOne platform. **(a)** Confusion matrix of the ResNet18-based binary classifier. On a manually labeled test dataset (n = 13,011 patches), the model achieved 1,056 true positives, 10,789 true negatives, 611 false negatives, and 265 false positives. **(b)** Receiver operating characteristic (ROC) curve of the classifier. The area under the curve (AUC) was 0.936, indicating strong discriminative power between true cells and background signals. **(c)** Representative examples of final cell detection results in dense and sparse regions. Columns show MIP images, detected cell centers, and composite overlays. White dots indicate automatically detected cell centers after blob detection and classification. Composite images demonstrate accurate localization of detected cells across tdTomato and eGFP channels, as well as in merged views. In the merged images, blue dots represent co-positive cells identified from blobs that show signals across multiple channels. **(d)** Region-wise quantification of normalized cell density across the ipsilateral hemisphere. Heatmaps represent region-wise cell density across multiple samples and injected regions. The analysis reveals distinct spatial patterns of cell distribution across brain regions, demonstrating the platform’s ability to integrate cell detection with atlas-based mapping and quantitative regional analysis.

The confusion matrix showed that, out of 13,011 test patches, the model correctly classified 1,056 true positives and 10,789 true negatives, with 265 false positives and 611 false negatives. From these results, the following performance metrics were computed: Area Under the Curve (AUC): 0.936, Accuracy: 93.6%, Specificity: 97.6%, Precision: 79.9%, Recall: 63.3%, and F1-score: 70.6%. These results demonstrate that the model maintains robust performance across a wide range of imaging conditions and anatomical variations. The consistently strong performance suggests that the classifier generalizes well and can be applied to heterogeneous brain imaging datasets without requiring extensive retraining. Notably, the model achieves relatively high precision with moderate recall, reflecting a conservative detection strategy that prioritizes minimizing false positives in large-scale datasets.

Visual inspection of prediction results on individual mouse brain samples confirmed that the model exhibited consistent performance across both high-density and low-density regions **(Fig. 3c**, **Fig. 4)**. For instance, in source regions with strong fluorescent signals, where viral injection led to dense cell labeling, the model accurately detected individual cells without over-segmentation, despite the high cellular density. Conversely, in more distal recipient regions with sparse cell distribution and weaker signal intensity, the model successfully detected true cells without omission, demonstrating robust sensitivity.

**Fig. 4.**
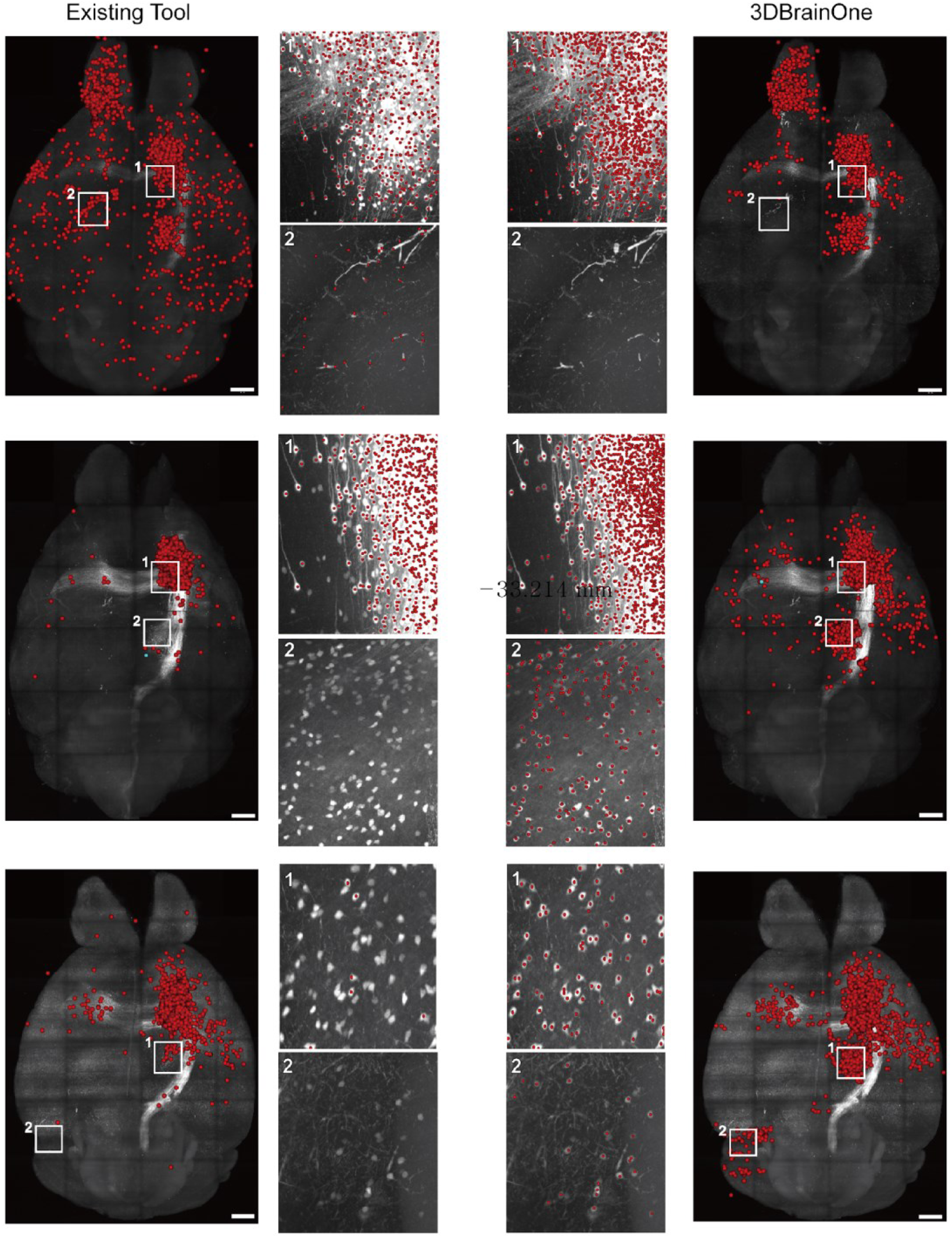
Qualitative comparison of cell detection results between an existing tool and 3DBrainOne across representative brain regions. Each row shows a different whole-brain sample, with red circles marking detected cells overlaid on 3D light-sheet fluorescence images. The left columns display results obtained from an existing detection tool in the eFLASH [24], while the right columns show outputs from 3DBrainOne on the same datasets. Zoomed-in panels (1, 2) highlight detection performance in both densely labeled and sparsely labeled regions. The existing tool often misses cells in dim areas (false negatives) or overdetects in noisy regions (false positives), especially in low-contrast zones. In contrast, 3DBrainOne achieves more accurate and anatomically consistent cell detection by using a permissive threshold combined with deep learning–based filtering, reducing false positives while preserving true cells.

This consistency across samples, despite variability in fluorescence intensity, tissue clearing quality, and imaging system configurations, can be attributed to several key design principles integrated into the platform. Specifically, by employing threshold-based blob detection rather than traditional segmentation, the model avoided artifacts from overlapping signals and remained sensitive to low-intensity signals. In addition, the training dataset was constructed by dividing each whole brain into a 5×5×5 grid and uniformly sampling patches from all subregions. This ensured that the model was exposed to both densely and sparsely populated regions during training, mitigating any bias toward high-density areas. Furthermore, the model was trained on a diverse collection of 21 whole-brain samples acquired under various imaging conditions, thereby enhancing its generalizability across heterogeneous experimental settings.

Together, these multi-layered strategies enabled the model to learn the structural heterogeneity of the whole brain without overfitting to any specific region or condition, ultimately achieving both high prediction accuracy and reproducibility across a wide range of biological and technical scenarios.

### Region-Wise Analysis Reveals Distinct Patterns of Interregional Signal Propagation

To further investigate the spatial organization of signal propagation across the brain, we performed region-wise analyses using normalized cell density data derived from the 3DBrainOne cell-counting module. By aggregating the number of detected cells in each anatomical region and normalizing by region volume, we visualized interregional variation as a heatmap **(Fig. 3d)**. The heatmap reveals heterogeneous patterns of signal distribution across brain regions, with a subset of regions exhibiting relatively elevated normalized densities relative to the global background.

Notably, the observed patterns vary among the injected sources (A–D), indicating that the distribution of the downstream signal is influenced by the source of the input. While most regions show low baseline levels, a limited number consistently exhibit higher signal enrichment, forming localized hotspots within the ipsilateral hemisphere. As these measurements are directly derived from cell-level detections, the observed regional differences reflect discrete cellular propagation patterns. This region-wise heterogeneity highlights that signal propagation is structured rather than random, and that distinct spatial patterns can be resolved through normalized density-based quantification.

### Voxel-based quantification enables region-specific analysis of brain volume

To enable regional-level structural analysis, we developed a voxel-based volumetric quantification module that estimates anatomical size directly from whole-brain 3D imaging data. This approach defines regional size by the number of voxels assigned to each anatomical region after atlas registration. When a minimal threshold (e.g., threshold = 1) was applied, the analysis included nearly all voxels within the brain volume. Under this condition, the voxel count per region serves as a direct proxy for anatomical volume, enabling consistent comparisons both within and across individual samples.

To ensure computational scalability for whole-brain datasets, voxel processing was batched during coordinate transformation, preventing excessive memory usage when handling large voxel sets. Image data were processed slice-by-slice rather than loading the entire volume into memory, further improving memory efficiency for large-scale datasets. Optional spatial subsampling along the xy plane and z-axis was also incorporated to accelerate processing, with voxel counts rescaled according to the sampling factor to approximate full-resolution measurements while maintaining comparability across samples.

For quality control and interpretability, the platform also provides native-space MIP visualization of selected voxels. In addition to global visualization, users can specify target anatomical regions; in this case, only voxels assigned to the selected regions are used to generate region-specific MIP images. This enables intuitive verification of voxel distribution and region assignment prior to downstream quantitative analysis.

### Region-Wise Volume Analysis Reveals Consistent Anatomical Scaling Across Brain Regions

To investigate structural variation across brain regions, we quantified region-specific volumetric profiles from voxel distributions. Across the whole brain, substantial regional volume differences were observed, reflecting known anatomical hierarchies **(Fig. 5a, 5b)**. Large structures, such as the cortical plate, exhibited markedly higher voxel counts than subcortical regions, including the thalamus and striatum, confirming that the platform accurately captures global brain organization.

**Fig. 5.**
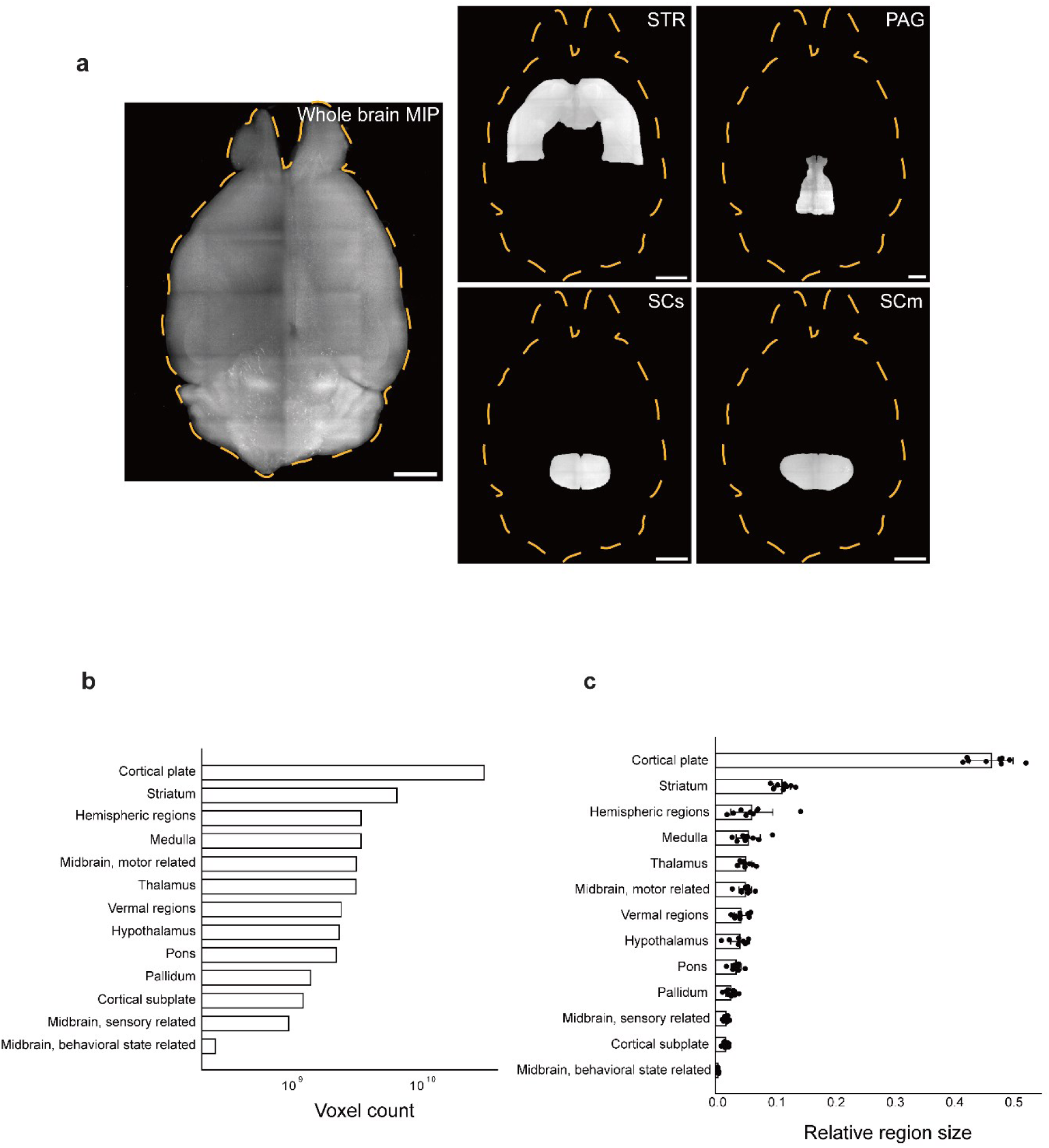
Voxel-based region-wise volume quantification and cross-sample consistency. **(a)** Validation of voxel-to-region assignment using MIP images. Representative atlas-mapped voxel distributions are shown for selected brain regions, including the striatum (STR), periaqueductal gray (PAG), superior colliculus sensory layer (SCs), and superior colliculus motor layer (SCm). Region-specific voxels are visualized after atlas registration, confirming accurate spatial localization and anatomical correspondence of assigned voxels. **(b)** Region-wise volumetric profile within a single brain sample. Regional size was quantified based on voxel counts assigned to each anatomical region. The resulting distribution reflects known anatomical hierarchies, with larger structures, such as the cortical plate, exhibiting substantially higher voxel counts than subcortical regions. **(c)** Cross-sample consistency of region-wise volume measurements. Relative region size was computed for nine independent brain samples by normalizing voxel counts to the total brain volume (sum of cerebrum, cerebellum, and brain stem). Mean values (bars) and individual sample measurements (dots) are shown for each region. Major anatomical regions exhibit consistent relative proportions across samples, demonstrating the robustness and reproducibility of voxel-based volumetric quantification.

Region-wise volume measurements were consistent across samples, with major anatomical structures maintaining similar relative proportions **(Fig. 5c)**. This indicates that the voxel-based quantification module preserves structural integrity and enables reliable comparison of anatomical size across datasets.

Notably, this approach provides a direct, scalable method for estimating region-wise anatomical size without requiring additional segmentation steps. The resulting volumetric profiles provide a robust structural reference to support downstream analyses of spatial distribution and regional variation in whole-brain imaging data.

## Discussion

In this study, we developed 3DBrainOne, an integrated platform designed for the 3D whole-brain single-cell-level image analysis. 3DBrainOne consolidates multi-step workflows—from cell detection and classification to atlas registration and quantification—into a unified, accessible environment.

While several tools currently exist for whole-brain imaging analysis, they often remain fragmented or require substantial technical expertise. For instance, Cellfinder [26], ClearMap [12] and the Chung lab workflow [24] provide robust cell counting and registration but rely on complex configurations and scripting, posing challenges for users without a computational background. Furthermore, these tools often lack functionality for multi-channel co-positivity analysis, manual refinement, and visual inspection to validate results. Other established tools, such as Brainreg [13] and Neuroglancer [14], excel at specific tasks, such as registration and 3D visualization, but do not offer a complete, all-in-one solution for automated quantification. Also, those tools do not have software for volumetric analysis. 3DBrainOne addresses these collective limitations by providing a streamlined, GUI-based environment that incorporates automated processing with the flexibility of manual modification. This design allows researchers to incorporate domain knowledge through iterative refinement without requiring computational expertise.

To ensure reliable results, the platform combines threshold-based blob detection with a deep-learning classifier, effectively balancing sensitivity to weak signals with the suppression of false positives. By employing uniform sampling across diverse brain regions during training, the system ensures consistent performance across heterogeneous anatomical contexts. Furthermore, the GUI-based implementation allows iterative calibration of detection parameters and model output thresholds, enabling users to interactively validate and refine results at each stage of the workflow. This allows users to validate and adjust results across varying signal densities and imaging conditions.

Integrating region-wise measurements with normalized cell density demonstrates our platform’s utility for identifying complex spatial trends in the brain. By quantifying interregional variations, the platform enables the detection of heterogeneous signal distributions that might otherwise be overlooked. These results demonstrate that our approach can resolve distinct spatial patterns across the brain.

An important extension of this platform is the incorporation of voxel-based region-wise volumetric quantification. By assigning atlas-registered voxels directly to anatomical regions, the platform enables scalable and consistent estimation of regional size. The resulting volumetric profiles not only recapitulate the expected hierarchical differences among major brain structures but also exhibit high consistency across independent samples. This consistency, particularly after normalization to total brain volume, supports reliable cross-sample comparison.

The versatility of 3DBrainOne extends beyond the specific applications presented in this study, as the platform is readily applicable to other modalities—such as transcriptomic mapping and viral tracing—where precise spatial quantification is critical. Its architecture is specifically designed to support the future integration of marker-specific classification models, allowing users to adapt the tool to various fluorescent reporters and experimental designs. While the current platform excels at robust binary classification, a remaining challenge is addressing increasingly complex cellular phenotypes that require multi-class identification. As imaging datasets scale in volume and variability, enhancing the platform’s computational efficiency and incorporating automated parameter optimization will complement the existing manual refinement capabilities. Future iterations will focus on these technical expansions and on integrating complementary data modalities, such as spatial transcriptomics, to provide an even more comprehensive and high-throughput tool for advanced brain mapping.

In summary, 3DBrainOne contributes to the field by providing a scalable, intuitive solution that lowers technical barriers to quantitative mouse brain mapping These developments will support more comprehensive investigation of whole-brain organization in both physiological and pathological contexts.

## Data Availability

All raw data described in this study are available from the corresponding authors upon request.

## Acknowledgements

The authors thank the members of the NBML (Neurotechnique and Brain Mapping Laboratory) for helpful discussions.

## Funding Statements

Y.-G.P. discloses support for the research of this work from the Ministry of Science and ICT, Republic of Korea [grant number RS-2024-00439379] and the Ministry of Health and Welfare, Republic of Korea [grant number RS-2023-00265963].

## Competing interests

The authors declare no competing interests.

## Notes

### Competing Interest Statement

The authors have declared no competing interest.

## Reference

[1] Ertürk, Ali, et al. “Three-dimensional imaging of solvent-cleared organs using 3DISCO.” Nature protocols 7.11 (2012): 1983–1995.

[2] Susaki, Etsuo A., et al. “Whole-brain imaging with single-cell resolution using chemical cocktails and computational analysis.” Cell 157.3 (2014): 726–739.

[3] Gao, Ruixuan, et al. “Cortical column and whole-brain imaging with molecular contrast and nanoscale resolution.” Science 363.6424 (2019): eaau8302.

[4] Economo, Michael N., et al. “A platform for brain-wide imaging and reconstruction of individual neurons.” elife 5 (2016): e10566.

[5] Susaki, Etsuo A., et al. “Advanced CUBIC protocols for whole-brain and whole-body clearing and imaging.” Nature protocols 10.11 (2015): 1709–1727.

[6] Chung, Kwanghun, et al. “Structural and molecular interrogation of intact biological systems.” Nature 497.7449 (2013): 332–337.

[7] Hansen, Henrik H., Urmas Roostalu, and Jacob Hecksher-Sørensen. “Whole-brain three-dimensional imaging for quantification of drug targets and treatment effects in mouse models of neurodegenerative diseases.” Neural Regeneration Research 15.12 (2020): 2255–2257.

[8] Dodt, Hans-Ulrich, et al. “Ultramicroscopy: three-dimensional visualization of neuronal networks in the whole mouse brain.” Nature methods 4.4 (2007): 331–336.

[9] Schindelin, Johannes, et al. “Fiji: an open-source platform for biological-image analysis.” Nature methods 9.7 (2012): 676–682.

[10] McQuin, Claire, et al. “CellProfiler 3.0: Next-generation image processing for biology.” PLoS biology 16.7 (2018): e2005970.

[11] Bankhead, Peter, et al. “QuPath: Open source software for digital pathology image analysis.” Scientific reports 7.1 (2017): 1–7.

[12] Renier, Nicolas, et al. “Mapping of brain activity by automated volume analysis of immediate early genes.” Cell 165.7 (2016): 1789–1802.

[13] Tyson, Adam L., et al. “Accurate determination of marker location within whole-brain microscopy images.” Scientific reports 12.1 (2022): 867.

[14] Google Research. Neuroglancer. https://zenodo.org/record/5573294

[15] Lindeberg, Tony. “Feature detection with automatic scale selection.” International journal of computer vision 30.2 (1998): 79–116.

[16] He, Kaiming, et al. “Deep residual learning for image recognition.” Proceedings of the IEEE conference on computer vision and pattern recognition. 2016.

[17] Wang, Quanxin, et al. “The Allen mouse brain common coordinate framework: a 3D reference atlas.” Cell 181.4 (2020): 936–953.

[18] Madisen, Linda, et al. “A robust and high-throughput Cre reporting and characterization system for the whole mouse brain.” Nature neuroscience 13.1 (2010): 133–140.

[19] Kaplitt, Michael G., et al. “Long-term gene expression and phenotypic correction using adeno-associated virus vectors in the mammalian brain.” Nature genetics 8.2 (1994): 148–154.

[20] Kügler, S., E. Kilic, and Mathias Bähr. “Human synapsin 1 gene promoter confers highly neuron-specific long-term transgene expression from an adenoviral vector in the adult rat brain depending on the transduced area.” Gene therapy 10.4 (2003): 337–347.

[21] Park, Young-Gyun, et al. “Protection of tissue physicochemical properties using polyfunctional crosslinkers.” Nature biotechnology 37.1 (2019): 73–83.

[22] Yaniv, Ziv, et al. “SimpleITK image-analysis notebooks: a collaborative environment for education and reproducible research.” Journal of digital imaging 31.3 (2018): 290–303.

[23] Fawcett, Tom. “An introduction to ROC analysis.” Pattern recognition letters 27.8 (2006): 861–874.

[24] Yun, Dae Hee, et al. “Uniform volumetric single-cell processing for organ-scale molecular phenotyping.” Nature Biotechnology 43.12 (2025): 2031–2042.

[25] Paszke, Adam, et al. “Pytorch: An imperative style, high-performance deep learning library.” Advances in neural information processing systems 32 (2019).

[26] Tyson, Adam L., et al. “A deep learning algorithm for 3D cell detection in whole mouse brain image datasets.” PLoS computational biology 17.5 (2021): e1009074.

